# Compositional Mediation Analysis for Microbiome Studies

**DOI:** 10.1101/149419

**Authors:** Michael B. Sohn, Hongzhe Li

**Affiliations:** Department of Biostatistics and Epidemiology, University of Pennsylvania

## Abstract

Motivated by recent advances in causal mediation analysis and problems in the analysis of microbiome data, we consider the setting where the effect of a treatment on an outcome is transmitted through perturbing the microbial communities or compositional mediators. Compositional and high-dimensional nature of such mediators makes the standard mediation analysis not directly applicable to our setting. We propose a sparse compositional mediation model that can be used to estimate the causal direct and indirect (or mediation) effects utilizing the algebra for compositional data in the simplex space. We also propose tests of total and component-wise mediation effects using bootstrap. We conduct extensive simulation studies to assess the performance of the proposed method and apply the method to a real metagenomic dataset to investigate the effect of fat intake on body mass index mediated through the gut microbiome composition.

## 1 Introduction

It has been shown that fat intake is associated with body mass index (BMI) (Bray and Popkin, 1998) and obesity is associated with the gut microbiome (Ley *et al.,* 2006; Turnbaugh *et al.,* 2006). From this information, a very natural question to ask is whether fat intake has some effects on BMI mediated through the perturbation of the gut microbiome. The approach to answering this type of questions is known as “mediation analysis”. Mediation analysis is a statistical method of studying the effect of a treatment or exposure on an outcome transmitted through intermediate variables, referred to as “mediators” or “intervening variables”. It has been widely applied in various disciplines such as sociology, psychology and epidemiology and become increasingly popular due to recent advances in causal inference (Pearl, 2000; Rubin, 2005; Imai *et al.,* 2010), which clarifies the assumptions needed for a causal interpretation. Until recently, mediation analysis has been restricted to a single mediator as depicted in Figure 1, and the effect of a treatment on an outcome transmitted through a mediator is often formulated and implemented within the framework of linear structural equation models (LSEMs).

**Figure 1:**
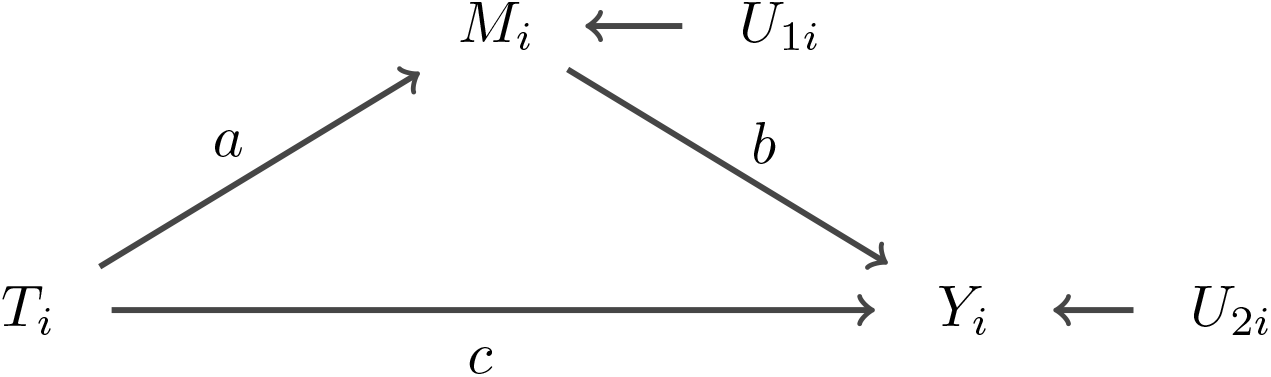
A single-mediator model: *a*, *b* and *c* are path coefficients; *U*_1*i*_ and *U*_2*i*_ are the disturbance or unmeasured variables for the mediator *M*_*i*_ and the outcome *Y*_*i*_, respectively.

For instance, an LSEM for the path diagram in Figure 1 can be formulated as

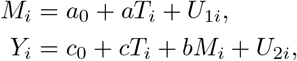

where *T*_*i*_ is a treatment, *M*_*i*_ a mediator and *Y*_*i*_ an outcome variable for unit *i*; *a*, *b* and *c* are path coefficients; *U*_1*i*_ and *U*_2*i*_ are disturbance variables for *M*_*i*_ and *Y*_*i*_, respectively. Under this model, the effect of *T*_*i*_ on *Y*_*i*_ transmitted through *M*_*i*_, often called the *indirect* or *mediation effect*, is generally defined by the product of the path coefficients *a* and *b*; the effect of *T*_*i*_ on *Y*_*i*_ not transmitted through *M*_*i*_, the *direct effect*, is defined by the path coefficient *c*. The model formulation for Figure 1 shows that the total effect of *T*_*i*_ on *Y*_*i*_ is the sum of the direct and indirect effects, *c + ab*.

In recent years, numerous studies have extended the applicability of mediation analysis: incorporating nonlinearity and interaction between the treatment and mediator (Pearl, 2001; Imai *et al*., 2010; VanderWeele and Vansteelandt, 2010); multiple mediators (Preacher and Hayes, 2008; VanderWeele and Vansteelandt, 2014). A few studies have also proposed methods for high-dimensional mediators. Chén *et al*. (2015) proposed a method to estimate path coefficients of an LSEM by finding the linear combinations of mediators that maximize the likelihood of linear structural equations, which is similar to the principal components. Huang and Pan (2016) introduced a transformation model using the principal components and included the interaction in their model. Zhao and Luo (2016) proposed a sparse mediation model using a regularized LSEM approach.

In this paper, we contribute to extending the applicability of mediation analysis further by proposing an estimating method for the causal direct and indirect effects when mediators are compositional. Compositional data refer to proportions or percentages of a whole and frequently arise in a wide range of disciplines such as mineral components of a rock in geology and vote shares of an election in psephology. In microbiome and metagenomic studies, to account for different sizes of sequencing libraries for 16S rRNA or shotgun metagenomic sequencing, the sequencing reads (i.e., count data) are often normalized into proportions. This normalization introduces the unit-sum constraint (i.e., proportions sum to unity), which transforms a *k* dimensional Euclidian space 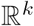 into a *k* −1 dimensional simplex space 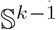, thus making the statistical models for unconstrained data inappropriate for compositional data. To deal with the nature of compositional data, Aitchison (1982) introduced an axiomatic approach with various operations under logratio transformation, which provides a one-to-one mapping between 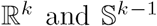, and various researchers including himself have formalized and extended this approach. Aitchison and Bacon-Shone (1984) proposed a linear and quadratic log contrast model for compositional covariates. Billheimer *et al*. (2001) formulated the algebra for compositions in the simplex space. Lin *et al*. (2014) developed a linear log contrast regression model for compositional covariates in a high-dimensional setting, and Shi *et al*. (2016) generalized the linear log contrast regression model.

We propose a framework for mediation analysis when mediators are high dimensional and compositional. Our framework utilizes two components: an estimation method based on the compositional operators of Aitchison (1982) and the composition algebra of Billheimer *et al*. (2001), and the linear log contrast regression of Lin *et al*. (2014); Shi *et al*. (2016). We employ the first component (i.e., compositional algebra) to jointly estimate the effect of a randomly assigned treatment on all compositional mediators. To this end, we propose to minimize the difference between observed and estimated compositions using the norm for a composition. We use the second component to quantify the effect of a treatment and compositional mediators on an outcome. Under this compositional mediation framework, we show that the causal direct and indirect effects are identifiable under some assumptions.

Section 2 introduces the compositional mediation model for continuous outcomes and discusses model assumptions and identifiability conditions. Section 3 describes methods of estimating composition and regression parameters and their covariance matrices. We also discuss null hypotheses for the total and component-wise mediation effects. Section 4 compares the performance of our method in extensive simulation studies with two methods that can be applied to compositional mediators. Section 4 also presents an application of the proposed method to a gut microbiome dataset. Finally, Section 5 presents a brief discussion of the methods and results.

## 2 Compositional Mediation Model and Causal Interpretation

### 2.1 Notation

For unit *i*, we let *T*_*i*_ be a treatment; ***M***_*i*_ be a vector of k compositional mediators; *Y*_*i*_ be an outcome; ***X***_*i*_ be a vector of *n*_*x*_ covariates that are nondescendants of *T*_*i*_ or ***M***_*i*_; ***U***_1*i*_ be a vector of disturbance variables for ***M***_*i*_; *U*_2*i*_ be a disturbance variable for *Y*_*i*_. We use 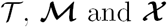 to denote the support of the distribution of *T*_*i*_, ***M***_*i*_ and ***X***_*i*_, respectively.

In the model assumptions and the identification of the causal direct and indirect effects, we adopt the potential outcomes framework. Let *M*_*i*_(*t*) denote the potential outcome for unit i under *T*_*i*_ = *t*, and *Y*_*i*_(*t,**m***) denote the potential outcome for unit *i* under *T*_*i*_ = *t* and ***M***_*i*_ = ***m***. Thus, we can express an observed variable ***M***_*i*_ = ***M***_*i*_(*T*_*i*_). Similarly, *Y*_*i*_ = *Y*_*i*_(*T*_*i*_,*M*_*i*_(*T*_*i*_)).

### 2.2 Compositional Mediation Model

Suppose we have a random sample of size n from a population where for each unit i we observe *Y*_*i*_, *T*_*i*_, and *M*_*i*_. For simplicity of notation, we assume ***X***_*i*_ = *X*_*i*_ is a one-dimensional covariate in the model formulation. See Supplementary Materials III for the model with multiple covariates ***X***_*i*_. Note that 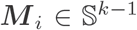 for all *i,* that is, 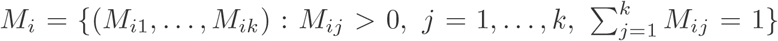. Figure 2 shows the effect of *T*_*i*_ on *Y*_*i*_ mediated through *M*_*i*_.

**Figure 2:**
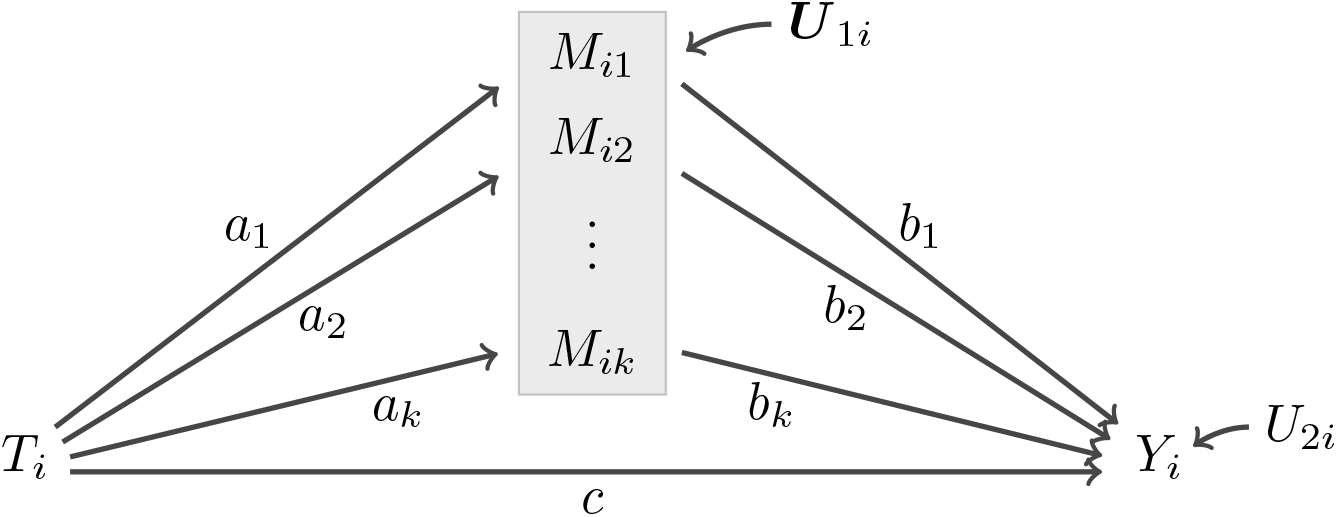
A compositional mediation model: *a*_*j*_, *b*_*j*_ and *c* are path coefficients, *j =* 1,…, *k*; *U*_1*i*_ and *U*_2*i*_ are disturbance variables for *k* compositional mediators *M*_*i*_ and an outcome *Y*_*i*_, respectively. 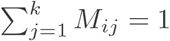.

Before introducing the compositional mediation model, we define two compositional operators as in Aitchison (1982). For two compositions 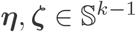, the perturbation operator is defined by

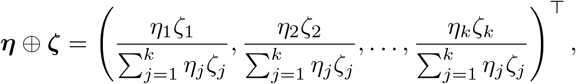

and the power transformation for a composition ***η*** by a scalar *α* by

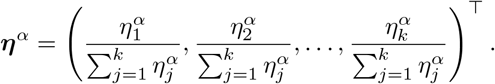

In addition, for a composition vector ***η***, define ***θ*** as the additive logratio transformation (alt) of ***η***, that is,

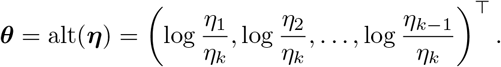

With these operators, we propose the following compositional mediation model:

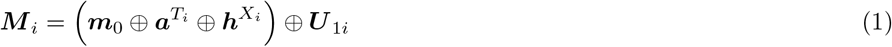

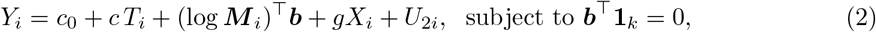

where ***m***_0_ is the baseline composition (i.e., when 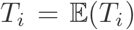); similarly, *c*_0_ is the baseline for *Y_i_, **a***, ***b*** and *c* are path coefficients; ***h*** and *g* are nuisance coefficients corresponding to the covariate *X*; **1**_*k*_ is a vector of *k* ones. We do not specify the distribution of *U*_1*i*_, but we assume *U*_2*i*_ ~ *N*(0, *σ*^2^). The model (1) formulates how a treatment perturbs a composition from the baseline composition, which is measured by the composition parameter *a.* With the compositional operators, all the calculations are within the simplex space, which leads to intuitive interpretation. In this model, the composition parameter *a* is directly interpretable as a composition, and a new composition is the baseline composition (*m*_0_) perturbed by *a* for *T*_*i*_ = 1. By taking the alt on both sides of model (1), we have

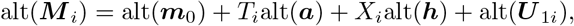

thus, alt(*U*_1*i*_) is modeled linearly with other terms.

The model (2) links a treatment and a composition to an outcome. To account for the compositional nature of *M*_*i*_, we impose a linear constraint, 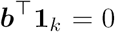, which is crucial for the estimator of regression coefficients to have desirable properties for compositional data, such as subcompositional coherence (Lin *et al.,* 2014; Shi *et al.,* 2016) since the regression coefficient *b* is scale-invariant with respective to *M*_*i*_, i.e., 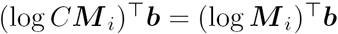 for any constant *C.*

Under this model, the total effect of *T*_*i*_ on *Y*_*i*_ can be decomposed into the direct effect, *c*, and the total indirect effect, 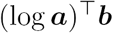, which is the sum of the component-wise indirect effects. Note that under the model (1), ***a*** is interpreted as the expected change in *M*_*i*_ due to *T*_*i*_ from the identity element 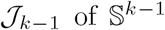, which is defined by 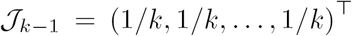. Thus, to estimate the component-wise indirect effects, ***a*** should be divided by 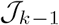, or multiplied by *k*, so that log(*ka*_*j*_) represents the expected change in log(*M*_*ij*_) from 0 by a one unit change in *T*_*i*_.

### 2.3 Model Assumptions and Identification

Identification of the causal direct and indirect effects requires several assumptions. To define our identifying assumptions, we use the potential outcomes notation described in Section 2.1. Combined with the stable unit treatment value assumption (SUTVA) and the positivity assumption (i.e., 0 < 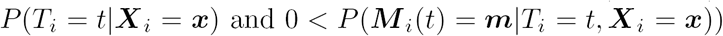, the assumptions for the compositional mediation model are given by

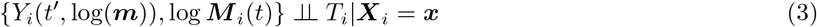

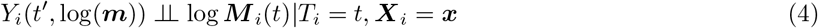

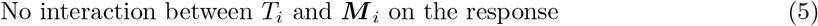

for 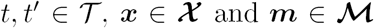. The SUTVA consists of two components: no interference (i.e., no effect of a treatment applied to one unit on an outcome for other units) and no hidden variation of treatment (i.e., consistent treatment levels) (Imbens and Rubin, 2015). Note that the second component is irrelevant for continuous treatments. The assumptions (3) - (4) with the positivity assumption are an extension of the sequential ignorability assumptions for the single mediator model (Imai *et al.,* 2010). Basically, they state that there is no unmeasured confounding variable after controlling covariates *X*_*i*_. The no interaction assumption (5) can be relaxed; See Supplementary Materials VI.

These assumptions appear like those for the multiple mediators model (Imai and Yamamoto, 2013; VanderWeele and Vansteelandt, 2014). However, in microbiome studies, the setting we consider is different from the multiple causal mechanisms with multiple mediators considered by Imai and Yamamoto (2013); VanderWeele and Vansteelandt (2014). We assume that the treatment *T* simply shifts the overall microbiome composition from one point in the simplex to another point, and therefore, all the components of a composition must be under the same treatment (i.e., log *M*_*i*_(*t*) or log *M*_*i*_(*t*′)); thus, there are, regardless of *k*, only four possible cases of conditional independence like the single mediator model,

i. 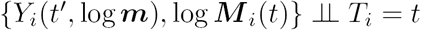
ii. 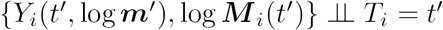
iii. 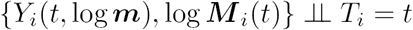
iv. 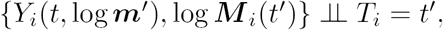

where log *m*′ is a vector of the potential values of log *M*_*i*_(*T*_*i*_) when *T_i_ = t′; t* is an observed treatment; *t'* is a reference value for the treatment. Similar conditions are required for the assumption (4).

Under the assumptions of the compositional mediation model, we have the following theorem to show that the model (1) and (2) lead to quantification of the causal direct and total indirect effects.

**Theorem 1** (Identification for the compositional mediation model) *Under the assumptions (3) - (5) combined with the SUTVA and the positivity assumptions, the causal direct effect ζ*(*t*) *and the causal total indirect effect σ*(*t*) *for the compositional mediation model are identifiable and given by*

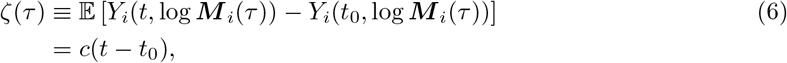

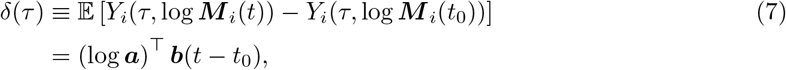

*where t is the observed treatment for unit i, t*_0_ *a reference value for the treatment and τ = t or t*_0_.

A proof of Theorem 1 is given in Supplementary Materials II. Pearl (2001) calls *ζ(τ)* the *average natural direct effect and Δ(δ)* the *average natural indirect effect.* Pearl (2001) also defines the *average controlled direct effect* that is defined in terms of a specific value of the mediator, rather than its potential values. Note that *ζ*(*t*) = ζ(*t*_0_) and *δ*(*t*) = δ(*t*_0_) under the no interaction assumption. Thus, the controlled and the natural direct effects coincide and are equal to c for a one unit change in *t*, and similarly, the natural total indirect effect is (log ***a***)^⊤^ ***b*** for a one unit change in *t.*

## 3 Parameter Estimation, Variance Estimation and Tests of Mediation Effects

### 3.1 Estimation of Composition Parameters and Covariance Matrix

With the composition operators, Billheimer *et al.* (2001) show that 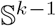 constitutes a complete inner product space, allowing the definition of a norm for a composition ***η***:

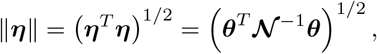

where *θ =* alt(η), and 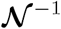 is the inverse matrix of a (*k* - 1) × (*k* - 1) matrix 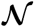 defined by

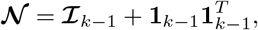

where 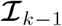 is the (*k* - 1) × (*k* - 1) identity matrix and 1_*k*-1_ is a (*k* - 1) column vectors of ones. Lastly, the inverse of the perturbation operator is defined by

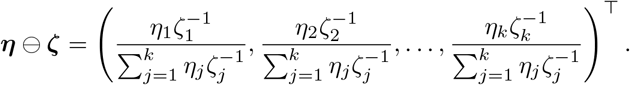

To estimate the parameters in the model (1), we propose the following objective function, which minimizes the composition norm of the difference between observed and estimated compositions,

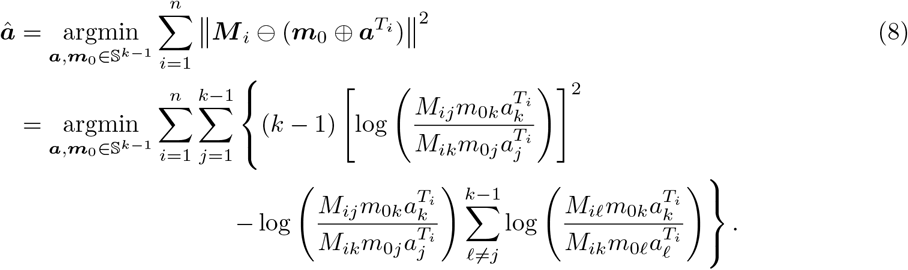

The objective function (8) is not convex in terms of *a*_*j*_, but is in terms of alt(*a*)_*j*_ for *j =* 1,…, *k* - 1. Thus, the optimal solution can be obtained by solving the following system of linear equations,

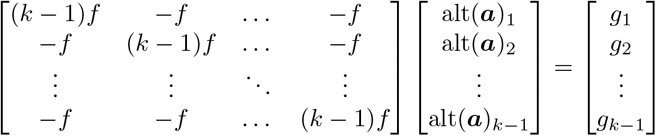

with a constraint 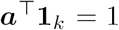, where 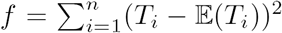 and 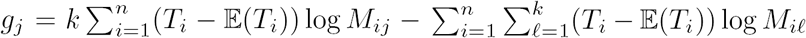. The matrix in the above equation is always invertible by Sherman-Morrison formula (Sherman and Morrison, 1950) since it can be expressed as the sum of a diagonal matrix of *k f* and a constant matrix of -*f*. Note that for brevity, we exclude *X*_*i*_; however, the presence of *X*_*i*_ does not fundamentally change the underlying method in estimating the composition parameters. For instance, if *n*_*x*_ covariates are present, the system of *n*_*x*_(*k* - 1) linear equations needs to be solved; See Supplementary Materials III for details. To estimate its covariance matrix 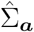, we use a bootstrap distribution of 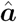 obtained by the percentile method of Machado and Parente (2005). See Supplementary Materials IV for details.

### 3.2 Estimation of Compositional Regression Parameters and Covariance Matrix

The log-contrast model has been the most general solution to incorporate the unit-sum constraint in the linear regression model for compositional covariates (Aitchison and Bacon-Shone, 1984; Lin *et al.,* 2014). Shi *et al.* (2016) developed a debias procedure for the *ℓ*_1_ regularized estimates of high dimensional compositional covariates. We use the linear log-contrast model and the debias procedure to estimate regression parameters, *b* and c, and their covariance matirx. Specifically, we first solve the following objective function,

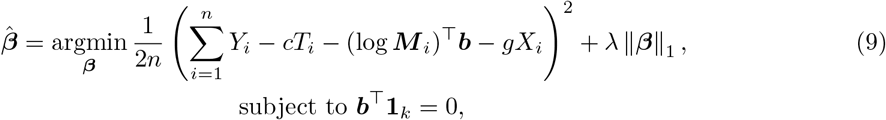

where *β =* (c, *b, g*)^⊤^ and λ is a turning parameter. For simplicity, the intercept is excluded in the model, which can be eliminated by centering all the variables in the model. We then apply the debias procedure of Shi *et al.* (2016) to the solution of the objective function (9) to obtain unbiased estimates and their covariance matrix.

### 3.3 Hypothesis Test of Mediation Effect

The null hypothesis of no total compositional mediation effect is given by

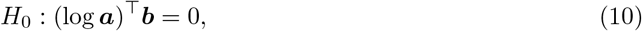

and the null hypothesis of no component-wise mediation effect is given by

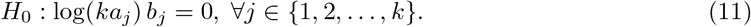

The null hypothesis (10) reflects the total mediation effect on an outcome; however, it can disguise the actual mediation effect, which may be captured by the null hypothesis (11). If for instance, the mediation effects of two mediators are equal, but their directions are opposite, then the total mediation effect is zero. In other words, we cannot reject the null hypothesis (10) but might reject the null hypothesis (11). Therefore, we need to test both hypotheses to avoid a misleading conclusion about the mediation effect.

To test the null hypotheses (10) and (11), we propose two approaches: an extension of the Sobel test (Sobel, 1982) and a bootstrap approach. In testing the null hypothesis (10) with the former, the square root of the first order asymptotic variance of a total indirect effect is computed with the estimated covariance matrices of 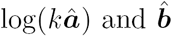 by the method described in Bollen (1987). It then is used as a standard error of the total indirect effect in the Z-test. The expressions for the first order asymptotic variances of the total indirect effect and component-wise indirect effects are given in Supplementary Materials V.

The distribution of a composition *M*_*i*_ is not known, but alt(*M*_*i*_)_*j*_ is well approximated by a normal distribution (Aitchison, 1986), that is, the distribution of log(*a*_*j*_/*a*_*k*_) is well approximated by a normal distribution. Therefore, *δ*(*τ*) can also be approximated by a normal distribution assuming the product of two normal variables (i.e., log(*a*_*j*_/*a*_*k*_) and *b*) follows a normal distribution. Recall that 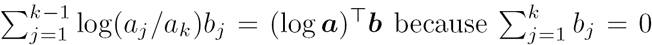. In general, the product of two normal variables does not follow a normal distribution. However, the misspecified distribution will just reduce the power but not affect the type I error rate when the null hypothesis of no indirect effect is false (MacKinnon *et al.,* 2002; Shrout and Bolger, 2002).

To avoid the assumption of normality for the indirect effect, we can use a bootstrap approach (Shrout and Bolger, 2002; VanderWeele and Vansteelandt, 2014). To this end, we use a non-parametric bootstrap for 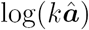 and a parametric bootstrap for 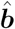 using a multivariate normal distribution to approximate the sampling distribution of *δ*(*τ*). The p-value for *δ*(*τ*) is then approximated by utilizing the fact that any bootstrap replicate *δ*(*τ*)_*b*_ - *δ*(*τ*) should have a distribution close to that of *δ*(*τ*) when the null hypothesis is true, where *δ*(*τ*)_*b*_ denotes an estimated total indirect effect derived from a resampled dataset (Efron and Tibshirani, 1993). That is, the p-value can be approximated by 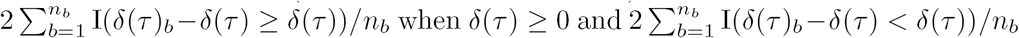 when *δ*(*τ*) < 0, where I(·) is the indicator function and *n*_*b*_ is the number of bootstrap samples. Similarly, the null hypothesis (11) can be tested. In the simulation, we observed no significant difference in performance between the two methods in terms of power and type I error, concurring with Huang and Pan (2016).

### 3.4 Sensitivity Analysis

In mediation analysis, the assumption (4) of no unmeasured confounding may not be satisfied even in a randomized experiment. In this case, the estimated total indirect effect is invalid since the parameter *b* in the model (2) is not consistent. To address this problem, we extend the single mediation model of Imai *et al.* (2010) to assess the sensitivity of an estimated total indirect effect to unmeasured pre-treatment confounding by utilizing the relationship among *U*_0*i*_, *U*_1*i*_ and *U*_2*i*_, where 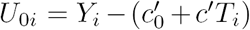 is the error in modeling the total treatment effect. Note that the total effect of a treatment is decomposed into the direct and total indirect effects under our model specification. Since we can treat *k* compositional mediators as a single mediator with *k* components, we assess the sensitivity of the proposed estimates by considering the correlation between the disturbance terms for the mediator and the outcome due to unmeasured confounding,

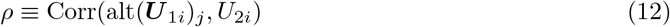

for all *j* = 1,…*k* – 1.

Suppose that the assumption (3) is satisfied and our model is correctly specified. Then, for a given correlation *ρ*, the causal total indirect effect is identified and given by

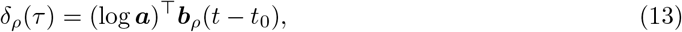

where *b*_*ρ*_ is the solution of the following system of *k* equations:

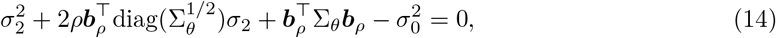

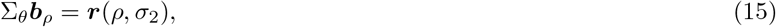

where 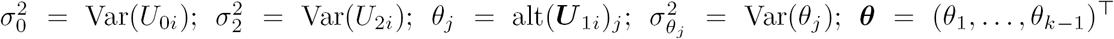; 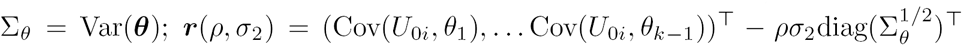. The derivation of the equations (14) and (15) is given in Supplementary Materials VII. Here, the parameters 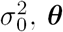, Σ_*θ*_ and Cov(*U*_0*i*_, θ_*j*_) for *j =* 1,…, *k* - 1 can be estimated consistently from the residuals, and 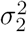 is a part of the solution to the above system of *k* equations.

We cannot extract any information about *ρ* from the data, we therefore treat *ρ* as a sensitivity parameter, choose a plausible range of *ρ*, and obtain corresponding values for TIDE.

## 4 Simulation Studies

Mediation analysis for multiple or high dimensional mediators typically assumes independence between mediators to establish a causal interpretation so principal components of mediators are often used. This approach of using principal components is also applicable for compositional mediators to estimate the direct and total indirect effects. Another naive approach for compositional mediators is to utilize an *ℓ*_1_ regularization, which tends to drop correlated variables. We used these two approaches to evaluate and compare the performance of our compositional mediation model. For the hypothesis test, we used the extension of the Sobel test for fair comparison.

In data generation, we randomly generated the treatment *T*_*i*_ from the standard normal distribution and the baseline composition *m*_0_ from the standard uniform distribution, Unif(0,1), under the unit-sum constraint. The path coefficients *a*, *b* and c were selected such that the direct effect is 1.00 and the total indirect effect is approximately 0.00, 0.50, 0.75 or 1.00. For the compositional disturbance, we used a multivariate logistic normal (LN) distribution (Aitchison, 1986) with mean 0_*k –* 1_ and covariance 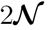, that is, 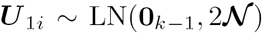. For the compositional regression disturbance, we used a normal distribution with mean 0 and variance 2, or *U*_2*i*_ ~ N(0, 2). Recall that our model does not specify the distribution of *U*_1*i*_. The composition *M*_*i*_ and the outcome *Y*_*i*_ were then generated according to the model (1) and (2), respectively.

### 4.1 Comparison of Power and Type I Error

We first estimated the power and type I error rate in testing the direct effect (DE) and the total indirect effect (TIDE) for three methods: the principal component regression (PCR), the two-stage adaptive lasso (TSAL) and our compositional mediation model (CMM). TSAL, as an *ℓ*_1_ regularization approach, uses the standard lasso in the first stage to screen irrelevant variables and the adaptive lasso in the second stage to select consistent variables (Bühlmann and van de Geer, 2011). We used 1500 simulations for each *k* mediators with a sample size *n =* 100 at various probability thresholds, where *k =* 5,49,99: 250 simulations with each of TIDE = 1.00,0.75,0.50; 250 simulations with no effect of *T*_*i*_ on *M*_*i*_ (i.e., *a_j_ =* 0, ∀*j*); 250 simulations with no effect of *M*_*i*_ on *Y*_*i*_ (i.e., *b_j_ =* 0, ∀*j*); 250 simulations with *inconsistent* TIDEs (i.e. the sum of the component-wise indirect effects is zero). For PCR, we included only the first *k*_*pc*_ principal components that explain 90% of the total variance. As shown in Table 1, while all three methods roughly control the type 1 errors, CMM outperforms both the PCR and TSAL approaches in power, especially when *k* is large.

To test the bias and variance of the estimates of the three methods, we simulated data with *c* = 1.00 and (log *a*)^*⊤*^*b =* 0.75 at various sample sizes *n =* 100, 200, 500, and the noises are generated from 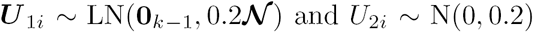. Figure 3 shows the results based on 200 simulations. When the sample size is small, the estimates of all three methods are slightly biased; however, as the sample size increases, they converge to the true values. The results are similar for *k =* 5,49 and 99.

**Table 1:** Power and type 1 error rate in testing TIDEs: number of mediators *k =* 5,49, 99; sample size *n =* 100; significance level *α =* 0.001,0.01,0.05. A total of 1500 simulations for each *k* were used: 750 simulations with non-zero TIDEs and 750 simulations for zero TIDEs. CMM: proposed compositional mediation model; PCR: principal component regression; TSAL: two-stage adaptive lasso.

|  |  | Power |  |  | Type 1 error |  |  |
| --- | --- | --- | --- | --- | --- | --- | --- |
| | $\alpha$ | 0.001 | 0.01 | 0.05 | 0.001 | 0.01 | 0.05 |
| $k = 5$ | CMM | 0.452 | 0.623 | 0.752 | 0.001 | 0.011 | 0.035 |
|  | PCR | 0.381 | 0.604 | 0.732 | 0.000 | 0.007 | 0.040 |
|  | TSAL | 0.413 | 0.621 | 0.740 | 0.001 | 0.008 | 0.037 |
| $k = 49$ | CMM | 0.476 | 0.675 | 0.820 | 0.001 | 0.015 | 0.043 |
|  | PCR | 0.051 | 0.207 | 0.423 | 0.000 | 0.005 | 0.029 |
|  | TSAL | 0.277 | 0.487 | 0.631 | 0.001 | 0.012 | 0.067 |
| $k = 99$ | CMM | 0.397 | 0.645 | 0.791 | 0.001 | 0.007 | 0.040 |
|  | PCR | 0.023 | 0.105 | 0.273 | 0.000 | 0.004 | 0.028 |
|  | TSAL | 0.304 | 0.495 | 0.628 | 0.003 | 0.016 | 0.064 |

**Figure 3:**
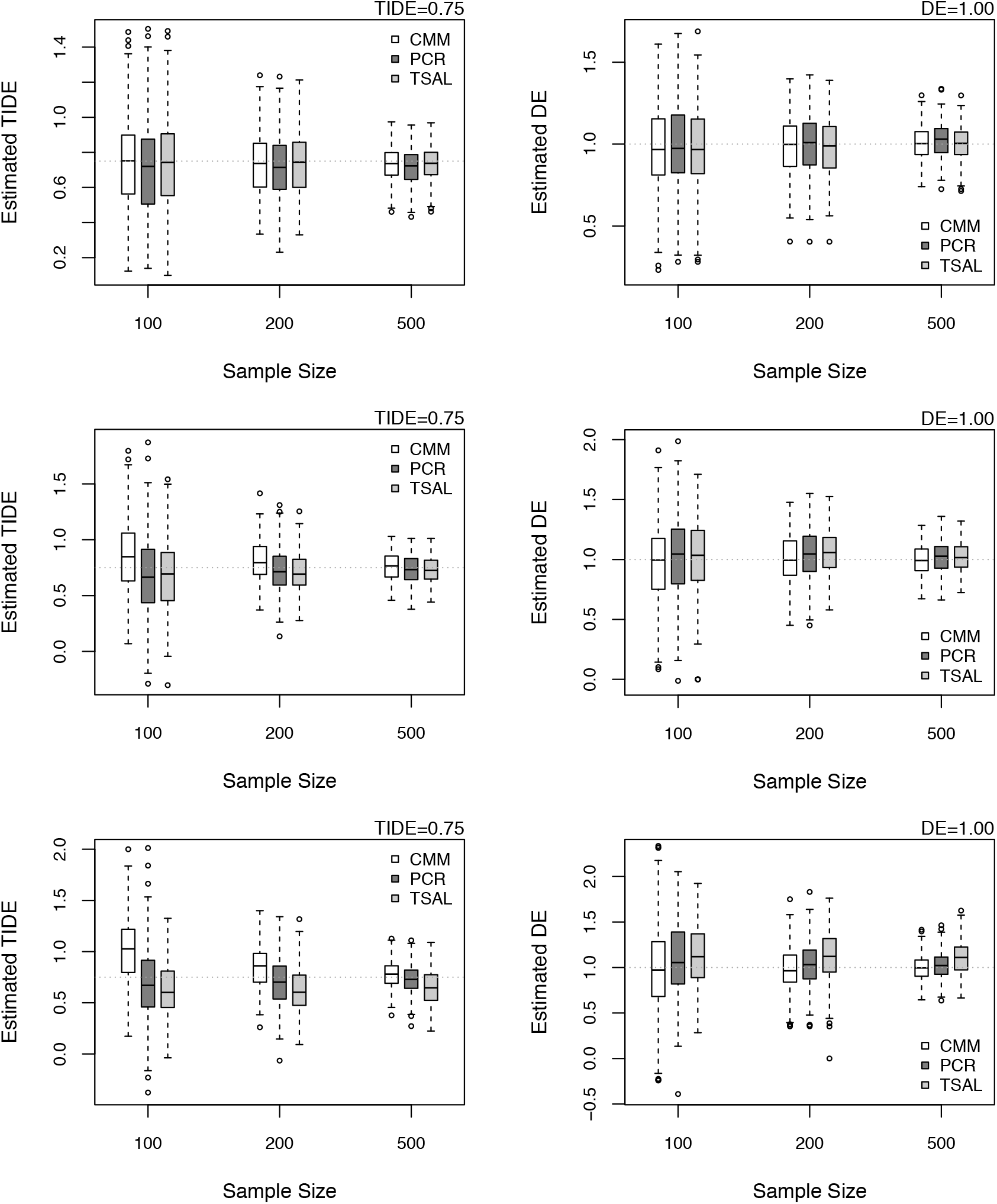
Estimated DEs and TIDEs at various sample sizes for the number of mediators *k* = 5 (top), *k* = 49 (middle) and *k* = 99 (bottom) with the true DE of 1.00 and the true TIDE of 0.75, which are indicated by dotted lines. The results are based on 200 simulations. CMM: proposed compositional mediation model; PCR: principal component regression; TSAL: two-stage adaptive lasso.

### 4.2 Identification of Component-wise Indirect Effect

PCR is not capable of testing the component-wise indirect effects (IDEs); therefore, we compared the performance of CMM on the component-wise IDEs only with TSAL. In data generation, we selected the parameters *a* and *b* such that the first 7 IDEs are at a nonzero constant level but different combinations of *a*_*j*_ and *b*_*j*_; the remaining 43 IDEs are at zero. The level of nonzero IDEs was increased by 0.1 from 0.0. We used sample sizes *n =* 100 and 200. For multiple testing corrections, we used the Benjamini and Yekutieli (2001) false discovery rate (FDR) at 0.05. As a comparison measure, the *F*_1_ score, which is the harmonic mean of precision and recall, was used. Figure 4 shows results: better performance of CMM over TSAL.

**Figure 4:**
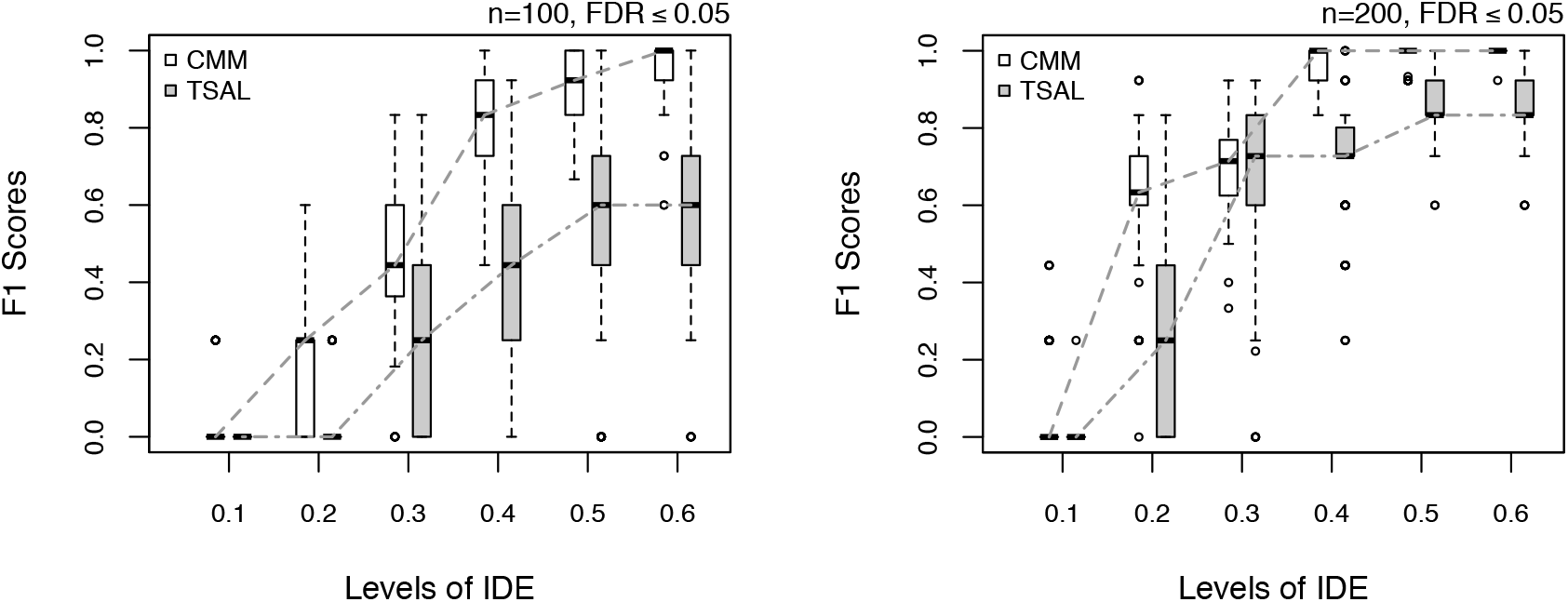
*F*_1_ score versus levels of IDEs at FDR ≤ 0.05. P-values are adjusted by the Benjamini-Yekutieli FDR procedure. CMM: proposed compositional mediation model; TSAL: two-stage adaptive lasso.

### 4.3 Simulation Based on Real Dataset

We also simulated data using the composition of a real dataset, COMBO, reported in Wu *et al.* (2011), which will be analyzed in Section 5. Specifically, the composition of *k =* 45 bacteria was available for a sample size of *n =* 98. The composition parameter *a* of this dataset was estimated by the Dirichlet regression (Maier, 2014) with randomly generated treatments *T*_*i*_ from N(0, 1). The regression parameter *b* under the linear constraint *b*^*⊤*^ 1_*k*_ = 0 was randomly generated from ±4 × Unif(0,1) for *k*_*s*_ components and set to 0 for *k*_*ns*_ components, where *k* = *k*_*s*_ + *k*_*ns*_, and *k*_*s*_ was randomly selected between 2 and 10. The direct effect c was set to 1, and we used *U*_2*i*_ ~ N(0, 2) to simulate the outcome *Y*_*i*_. Figure 5 shows that the estimates of DE and TIDE are almost unbiased for all three methods.

**Figure 5:**
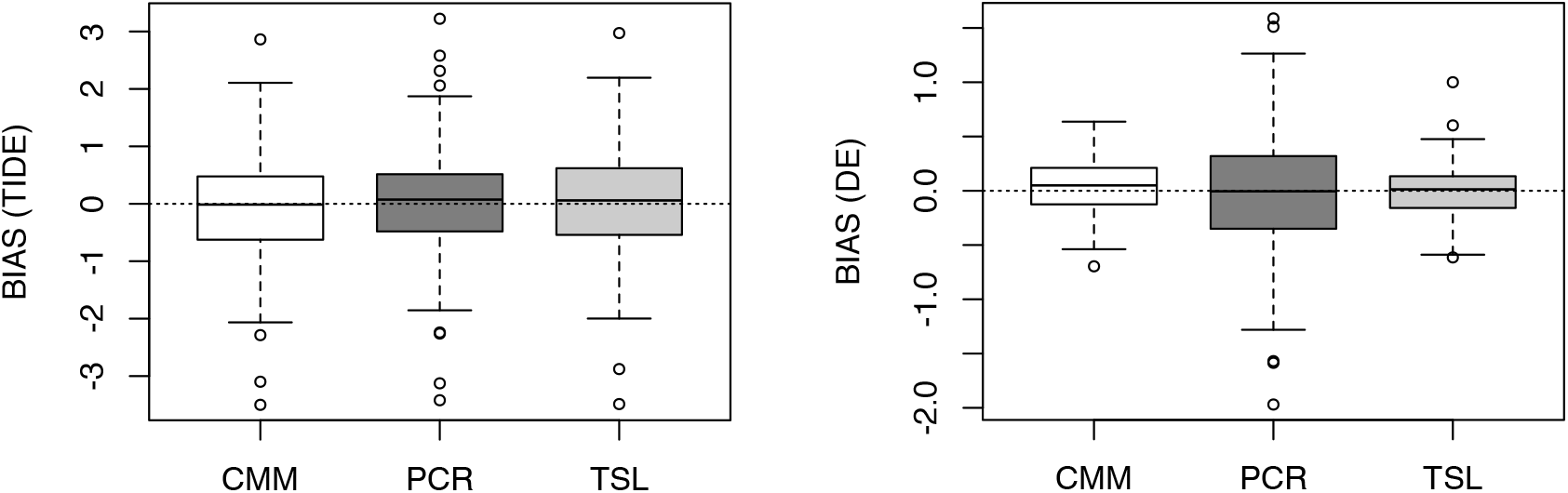
Biases of DEs and TIDEs for the data simulated with the composition of a real dataset. The results are based on 200 simulations. CMM: proposed compositional mediation model; PCR: principal component regression; TSAL: two-stage adaptive lasso.

Since we randomly generate TIDE, obtaining a particular value of TIDE is not computationally practical. Thus, we used (log *a*)^⊤^*b* > 0.75 to test the power. Based on 200 simulations, the power was 0.52, 0.38 and 0.48 respectively for CMM, PCR and TSAL for an *α* level of 0.05, and 0.24, 0.16 and 0.21 for an *α* level of 0.01. CMM still has the highest power while controlling the type I error.

## 5 Real Data Application

We applied CMM to a cross-sectional analysis of 98 healthy volunteers studied in the COMBO dataset (Wu *et al*., 2011), which is the source of motivation for this study. The dataset consists of 16S rRNA sequences from fecal samples of 98 healthy individuals from the University of Pennsylvania. It also contains demographic and clinical information including fat intake and BMI, where the habitual long-term fat intake were derived from the food frequency questionnaire (FFQ). Such measurements are widely applied in nutritional research, and their reproducibility and validity have been validated (Hu *et al*., 1999). We summarized operational taxonomic units (OTUs) at the genus level and then filtered out the genera that appear in fewer than 10% of the samples, leaving 45 genera in 98 samples. Due to different total counts throughout the samples, the OTU counts assigned to these genera were transformed into proportions after replacing zero counts by the maximum rounding error 0.5, which is commonly used in compositional data analysis (Aitchison, 1986).

### 5.1 Estimation of TIDE

The gut microbiota can influence host adiposity through energy extraction from the diet, with variable efficiency depending on community composition; furthermore, the microbiota can also affect host adiposity by influencing metabolism throughout the body. It is therefore highly likely that gut microbiome can potentially mediate the effect of the diet such as fat intake on host adiposity and BMI. Since the 98 samples were roughly randomly sampled, it is reasonable to assume that fat intake was randomly assigned. CMM was applied to the dataset with BMI as the outcome, fat intake as the treatment, and the 45 genera as the compositional mediators. The estimated DE and TIDE are 0.933 with a 95% CI of (0.005, 1.935) and 0.809 with a 95% CI of (-0.452, 2.250). The estimated component-wise IDEs are shown in Figure 6.

**Figure 6:**
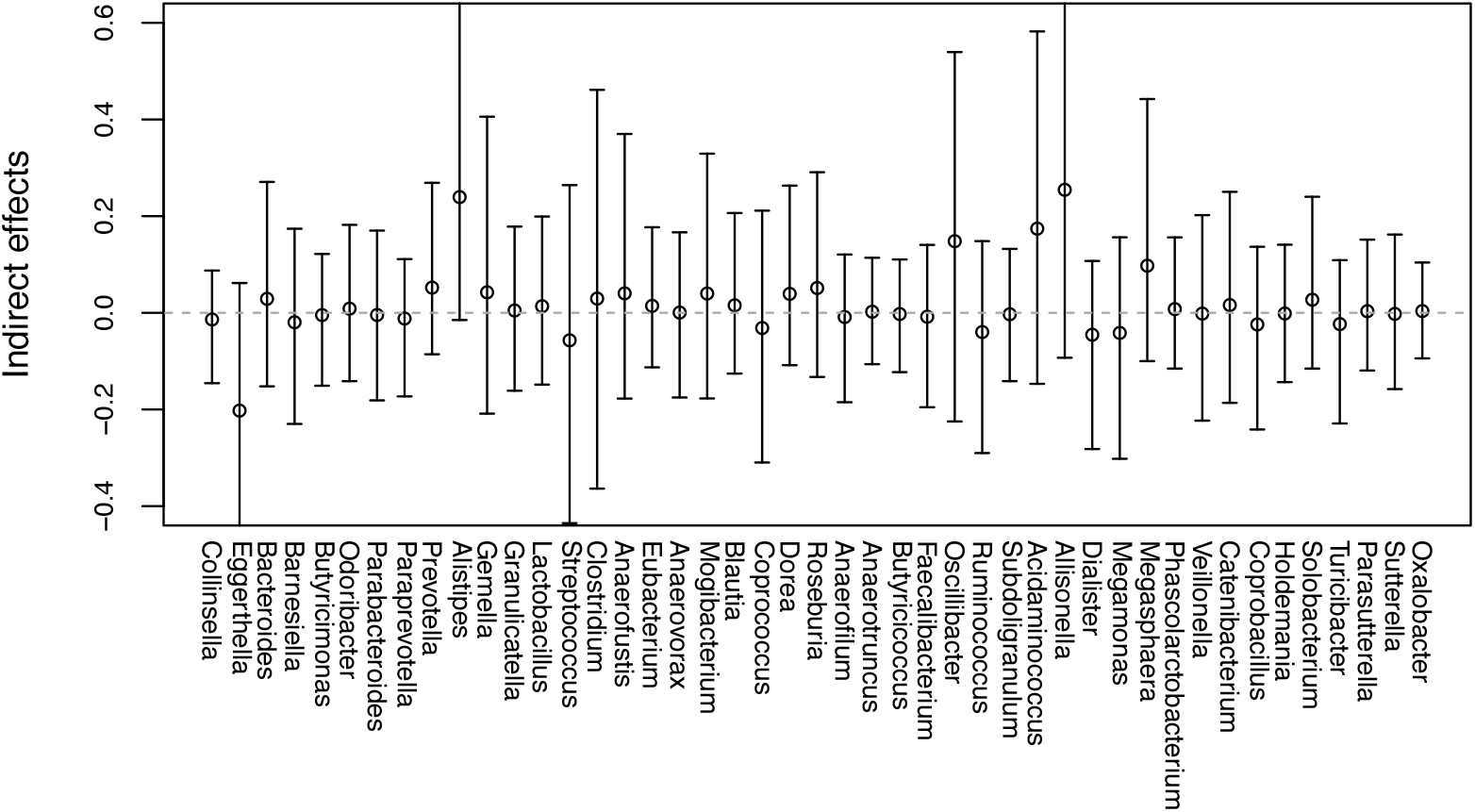
The estimated component-wise IDEs of fat intake on BMI through the gut microbiome. CI for *Eggerthella* is (-0.652, 0.062); CI for *Alistipes* is (-0.015, 0.668); CI for *Allisonella* is (-0.093, 0.786).

The estimated DE and TIDE (i.e., 0.933 and 0.808, respectively) correspond to 53.6% and 46.4% of the total effect of fat intake on BMI. Note that under our model specification, the total effect of a treatment can be decomposed into the direct and indirect effects. The estimated DE is statistically significant at the significance level of 0.05 but the estimated TIDE, as well as all the componentwise IDEs, are not. This insignificance of TIDE and some of the component-wise IDEs is likely due to insufficient sample size: based on the simulation study in which we used the characteristics of parameters estimated from the COMBO dataset, TIDE of 0.75 and component-wise IDEs at around 0.2 were rarely detected with a sample size of 100.

As shown in Figure 6, the potential genera that would have statistically significant non-zero component-wise IDEs with a sufficient sample size are *Eggerthella, Alistipes, Oscillibacter, Acidaminococcus* and *Allisonella*. All these genera except *Eggerthella* have positive values for IDEs, but their responses to fat intake are quite different. The abundance of *Alistipes* and *Oscillibacter* is negatively correlated with fat intake and BMI, whereas that of *Acidaminococcus* and *Allisonella* is positively correlated with fat intake and BMI. Based on this information, we can hypothesize that the increase in fat intake causes the decrease in the abundance of *Alistipes* and *Oscillibacter* and the increase in the abundance of *Acidaminococcus* and *Allisonella* which in turn cause the increase in BMI. Lam *et al*. (2012) identified Oscillibacter-like organisms as a potentially important gut microbe that mediates high fat-induced gut dysfunction and gut permeability and showed that decrease of *Oscillibacter* led to increased gut permeability, which was shown to be associated with obesity (Teixeira *et al*., 2012). This observation is largely consistent with those observed in mice fed with high-fat diet (Daniel *et al*., 2014).

As a comparison, TSAL gave the following estimates of DE = 0.751 and TIDE = 0.555, which correspond to 57.5% and 42.5% of the total effect of fat intake on BMI. TSAL selected five genera: *Alistipes* (0.161), *Clostridium* (0.031), *Doria* (0.047), *Acidaminococcus* (0.150), and *Allisonella* (0.166), where the values in parenthesis are the estimated component-wise indirect effects. All except DE were not significant at the significance level of 0.05, similar to the results of CMM. Interestingly, *Oscillibacter* was not selected by TSAL.

### 5.2 Sensitivity Analysis on COMBO

TIDE of fat intake on BMI is estimated under the assumptions (3) ~ (5). Even though we can assume random assignment of fat intake (i.e., the assumption (3) holds) as mentioned before, we cannot safely assume that the assumption (4) is satisfied. Thus, we estimated TIDE given the correlation *ρ* between the disturbance terms of the mediator and outcome using the method proposed in Section 3.4. Figure 7 shows this result. The estimated TIDE ranges from 2.162 to −1.546 for *ρ* ranges from −0.244 to 0.244. These values are within the 95% CI of the estimated TIDE assuming *ρ* = 0.

**Figure 7:**
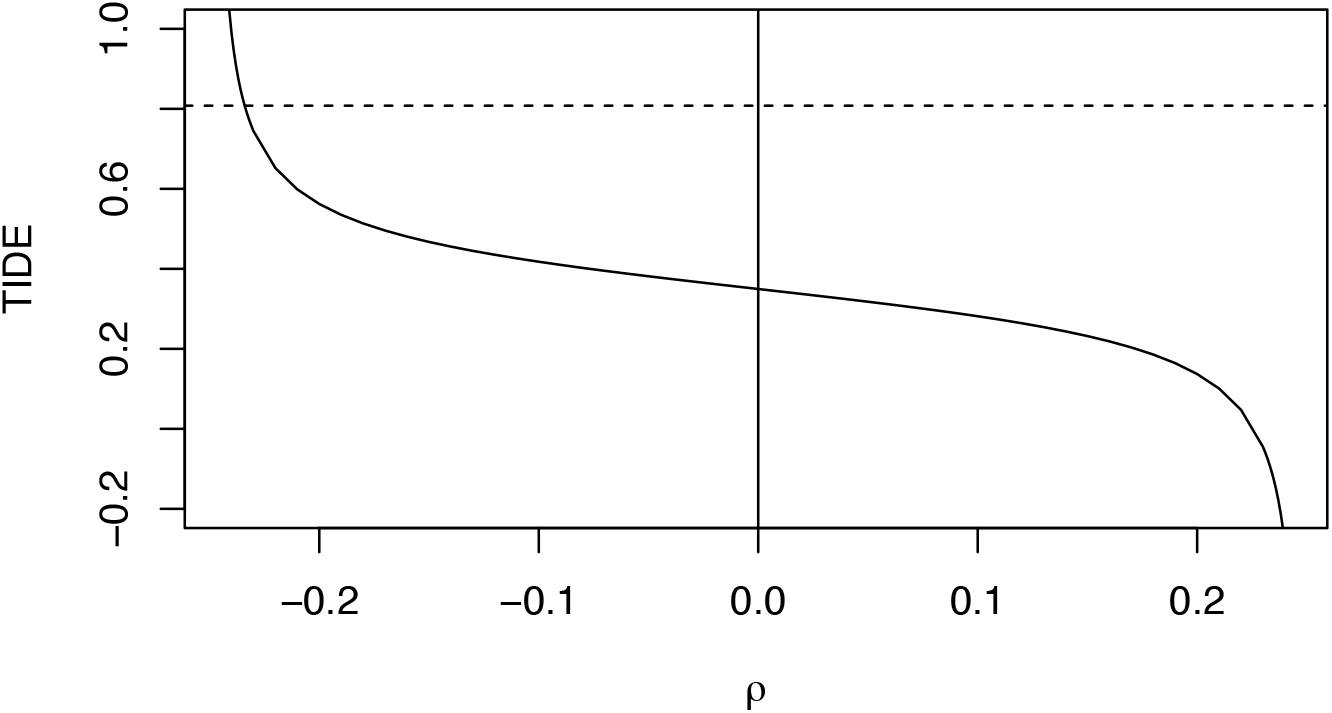
TIDE for given the correlations *ρ* between the disturbance terms of the mediator and outcome. The dashed line is the estimated TIDE (i.e., 0.808) by CMM. The solid line denotes TIDE for given *ρ.* The vertical line at 0 is the 95% bootstrap confidence interval (−1.735, 2.706) of TIDE when *ρ =* 0 (the *y*-axis is truncated at 1 for better visualization).

Our simulation results shown in Supplementary Materials VIII indicate that when the sample size is small, TIDE can be sensitive to the no confounding assumption. However, as the sample size increases,

## 6 Discussion

In this study, we have proposed a compositional mediation model for a continuous outcome. Our method takes the characteristics of compositional data into account and treats the whole compositional mediators as a unit, that is, it estimates the effect of a treatment on compositional mediators simultaneously instead of each mediator separately. In our simulation studies, we have shown better performance of our method over the two potential methods for compositional mediators. Our method also provides a clear interpretation of component-wise indirect effects. Application to the gut microbiome and BMI dataset has indicated potentially mediating effect of the gut microbiome in linking fat intake and BMI. Although the results were not statistically significant, several bacterial genera have been shown to be directly associated with gut permeability and therefore BMI. It would be interesting and important to replicate these results in larger data sets.

Even though we used a continuous treatment variable in the simulation studies, a binary treatment variable can be used without any modification to our method. In many clinical microbiome studies, the outcome variable is binary such as whether a subject is diseased or not. In these cases, Model (2) can be rewritten for logistic or probit regression, assuming the outcome variable is a latent continuous variable indicated by an observed dichotomous variable. Then, Model (1) and the modified Model (2) will provide the identifiability of the direct and indirect effects (Winship and Mare, 1983). Another interesting extension of our method is for longitudinal compositional data, which is also very common in microbial studies.

The Gaussian assumption is made for the error term in modeling the outcomes (BMI in our application), which often can be approximately achieved by an appropriate transformation. One important future research is to develop bootstrap methods for inference of DE and TIDE allowing heteroscedastic errors and heavy-tailed distributions.

## References

Aitchison, J. (1982). The statistical analysis of compositional data. J. R. Stat. Soc., Ser. B (Stat. Methodol.), 44(2), 139–177.

Aitchison, J. and Bacon-Shone, J. (1984). Log contrast models for experiments with mixtures. Biometrika, 71(2), 323–330.

Aitchison, J. (1986). The Statistical Analysis of Compositional Data. New York: Chapman & Hall.

Benjamini, Y. and Yekutieli, D. (2001). The Control of the False Discovery Rate in Multiple Testing Under Dependency. The Annals of Statistics, 29(4), 1165–1188.

Billheimer, D., Guttorp, P. and Fagan, W. F. (2001). Statistical Interpretation of Species Composition. Journal of the American Statistical Association, 96(456), 1205–1214.

Bollen, K. A. (1987). Total, direct, and indirect effects in structural equation models. Sociological Methodology. 17, 37–69.

Bray, G. A. and Popkin, B. M. (1998). Dietary fat intake does affect obesity! Am J Clin Nutr. 68(6), 1157–1173.

Bühlmann, P. and van de Geer, S. (2011). Statistics for High-Dimensional Data: Method, Theory and Applications. Berlin: Springer.

Chén, O. Y., Crainiceanu, C. M., Ogburn, E. L., Caffo, B. S., Wager, T. D. and Lindquist, M. A. (2015). High-dimensional Multivariate Mediation with Application to Neuroimaging Data. arXiv:1511.09354

Daniel, H., Gholami, A. M., Berry, D., Desmarchelier, C., Hahne, H., Loh, G., Mondot, S., Lepage, P., Rothballer, M., Walker, A., Böhm C., Wenning, M., Wagner, M., Blaut, M., Schmitt-Kopplin, P., Kuster, B., Haller, D. and Clavel, T. (2014). High-fat diet alters gut microbiota physiology in mice. ISEM J, 8(2), 295–308.

Efron, B. and Tibshirani, R. (1993). An Introduction to the Bootstrap. Chapman & Hall.

Hu, FB, Rimm, E., Smith-Warner, S.A., Feskanich, D., Stampfer, M.J., Ascherio, A., Sampson, L. and Willett, W.C. (1999). Reproducibility and validity of dietary patterns assessed with a food-frequency questionnaire. Am J Clin Nutr., 69(2), 243–249.

Huang, Y. T. and Pan, W. C. (2016). Hypothesis test of mediation effect in causal mediation model with high-dimensional continuous mediators. Biometrics, 72(2), 402–413

Lam, Y. Y., Ha, C. W., Campbell, C. R., Mitchell, A. J., Dinudom, A., Oscarsson, J., Cook, D. I., Hunt, N. H., Caterson, I. D., Holmes, A. J. and Storlien, L. H. (2012). Increased Gut Permeability and Microbiota Change Associate with Mesenteric Fat Inflammation and Metabolic Dysfunction in Diet-Induced Obese Mice. PLoS ONE, 7(3): e34233.

Imai, K., Keele, L. and Yamamoto, T. (2010). Identification, Inference and Sensitivity Analysis for Causal Mediation Effects. Statistical Science, 25(1), 51–71.

Imai, K., Keele, L. and Tingley, D. (2010). A General Approach to Causal Mediation Analysis. Psychol Methods, 15(4), 309–334.

Imai, K. and Yamamoto, T. (2013). Identification and Sensitivity Analysis for Multiple Causal Mechanisms: Revisiting Evidence from Framing Experiments. Political Analysis, 21(2), 141–171.

Imbens, G. and Rubin, D. (2015). Causal Inference for Statistics, Social, and Biomedical Sciences: An Introduction. Cambridge: Cambridge University Press.

Machado, J. A. F. and Parente, P. (2005). Bootstrap estimation of covariance matrices via the percentile method. The Econometrics Journal, 8(1), 70–78.

MacKinnon, D. P., Lockwood, C. M., Hoffman, J. M., West, S. G. and Sheets, V. (2002). A comparison of methods to test mediation and other intervening variable effects. Psychological Methods, 7(1), 83–104.

Maier, M. J. (2014). DirichletReg: Dirichlet Regression for Compositional Data in R. Research Report Series / Department of Statistics and Mathematics, 125. WU Vienna University of Economics and Business, Vienna.

Ley, R. E., Turnbaugh, P. J., Klein, S. and Gordon, J. I. (2006). Human gut microbes associated with obesity. Nature, 444, 1022–1023.

Lin, W., Shi, P., Feng, R. and Li, H. (2014). Variable selection in regression with compositional covariates. Biometrika, 101(4), 785–797.

Pearl, J. (2000). Causality: Models, Reasoning, and Inference. New York: Cambridge University Press.

Pearl, J. (2001). Direct and indirect effects. In Proceedings of the Seventeenth Conference on Uncertainty and Artificial Intelligence, 411–420. San Francisco, CA: Morgan Kaufmann.

Preacher, K. J. and Hayes, A. F. (2008). Asymptotic and resampling strategies for assessing and comparing indirect effects in multiple mediator models. Behav res methods, 40(3), 879–891.

Robins, J. M. (2003). Semantics of causal DAG models and the identification of direct and indirect effects. In Highly Structured Stochastic Systems (P. J. Green, N. L. Hjort and S. Richardson, eds.) 70–81. Oxford: Oxford Univ. Press.

Rubin D. B. (2005). Causal inference using potential outcomes. Journal of the American Statistical Association. 100 (469), 322–331.

Sherman, J. and Morrison, W. J. (1950). Adjustment of an Inverse Matrix Corresponding to a Change in One Element of a Given Matrix. Annals of Mathematical Statistics. 21(1), 124–127.

Shi, P., Zhang, A. and Li, H. (2016). Regression Analysis for Microbiome Compositional Data. Annals of Applied Statistics, 10(2), 1019–1040.

Shrout, P. E. and Bolger, N. (2002). Mediation in Experimental and Nonexperimental Studies: New Procedures and Recommendations. Psychological Methods, 7(4), 422–445.

Sobel, M. E. (1982). Asymptotic confidence intervals for indirect effects in structural equation models. Sociological methodology, 13, 290–312.

Teixeira, T. F., Collado, M. C., Ferreira, C. L., Bressan, J. and Peluzio, M. C. (2012). Potential mechanisms for the emerging link between obesity and increased intestinal permeability. Nutr Res., 32(9), 637–47.

Turnbaugh, P. J., Ley, R. E., Mahowald, M. A., Magrini, V., Mardis, E. R. and Gordon, J. I. (2006). An obesity-associated gut microbiome with increased capacity for energy harvest. Nature, 444, 1027–1031.

VanderWeele, T. J. and Vansteelandt, S. (2010). Odds ratios for mediation analysis for a dichotomous outcome. American Journal of Epidemiology 172(12), 1339–1348.

VanderWeele, T. J. and Vansteelandt, S. (2014). Mediation Analysis with Multiple Mediators. Epidemiol Method. 2(1): 95–115.

Winship, C. and Mare, R. D. (1983). Structural Equations and Path Analysis for Discrete Data. The American Journal of Sociology. 89(1):54–110.

Wu, G., Chen, J., Hoffmann, C., Bittinger, K., Chen, Y. Y., Keilbaugh, S. A., Bewtra, M., Knights, D., Walters, W. A., Knight, R., Sinha, R., Gilroy, E., Gupta, K., Baldassano, R., Nessel, L., Li, H., Bushman, F. D. and Lewis J. D. (2011). Linking long-term dietary patterns with gut microbial enterotypes. Science. 334(6052) 105–108.

Zhao, Y. and Luo, X. (2016). Pathway Lasso: Estimate and Select Sparse Mediation Pathways with High Dimensional Mediators. arXiv:1603.07749.

